# Individual differences in learning and decision-making: the role of COMT Val158Met polymorphism in transitive inference

**DOI:** 10.1101/2025.07.09.663879

**Authors:** Ann Paul, Mariella Segreti, Isabel Beatrice Marc, Maria Teresa Fiorenza, Sonia Canterini, Surabhi Ramawat, Giampiero Bardella, Pierpaolo Pani, Stefano Ferraina, Emiliano Brunamonti

## Abstract

Understanding the ordinal relationships between items requires constructing a rank order supporting decision-making between options. This process depends on the ability to learn reciprocal relationships and to select the best option available when making a choice. In such forms of decision-making, the prefrontal cortex (PFC) plays a crucial role in encoding the relative value of alternatives as a decision is formed. Higher-order cognitive abilities are influenced by genetic factors that affect dopamine availability in the PFC, potentially contributing to individual differences. Here, we examined the performance of 83 participants in a transitive inference task (TI), grouped by genotype based on the Val158Met single-nucleotide polymorphism in the *Catechol-O-Methyltransferase* (*COMT*) gene. The task included a learning phase in which participants acquired the reciprocal relationships among a set of hierarchically ranked items (A>B>C>D>E>F), followed by a test phase in which they were required to compare all possible item pairs and select the higher-ranked one. While genotype did not significantly influence test-phase performance, it did affect learning efficiency. Specifically, Val homozygotes took a longer learning procedure than both heterozygotes and Met homozygotes during the learning phase. Drift diffusion modelling (DDM) revealed that task performance was explained by the efficiency of evidence accumulation, which was lower in Val homozygotes, accounting for their poorer performance not only during initial learning but also when required to switch to a reversed hierarchical structure (A<B<C<D<E<F). These findings suggest that individual differences in inferential decision-making and cognitive flexibility may be partially driven by genetically determined variations in prefrontal dopamine availability.

## 1. Introduction

The ability to infer relationships between events from prior experiences is fundamental to both human and animal cognition (Bryant & Trabasso, 1971; Grosenick et al., 2007; Takahashi et al., 2008; Bond et al., 2010; Brunamonti et al., 2016). This ability minimises the amount of time spent making potentially dangerous trial-and-error decisions. Deducing that A>C after learning the independent premises A>B and B>C is a classic example. The cognitive mechanisms behind this type of reasoning, inferential reasoning, have been widely studied using the Transitive Inference (TI) task (Bryant & Trabasso, 1971; Ramawat et al., 2023). Participants in this task first learn the ordinal sequence of a set of ranked items (e.g., A>B>C>D>E>F) by identifying the reciprocal relationship between adjacent rank pairs (e.g., A>B) through trial-and-error. During a subsequent test phase, participants must respond to which items are higher in rank in all possible pairs, learned and novel (e.g., B>D). According to one of the effects observed during the TI test phase, the symbolic distance effect (SDE), comparisons between items with higher rank differences are generally faster and more accurate than those between items with smaller rank differences (Gazes et al., 2023). This effect is thought to rely on temporarily engaging a mental representation of their rank for the duration required to perform the comparison, where the degree of overlap in item representation interferes with the decision-making process (Ramawat et al., 2023). Studies have identified the involvement of a distributed network of brain regions during TI, with a central role played by frontal brain areas (Acuna D. et al., 2002; Brunamonti et al., 2016b; Mione et al., 2020; Ramawat et al., 2022; Zhang et al., 2022). Specifically, the prefrontal cortex (PFC) is known to encode abstract representations, maintaining the decision variables during deliberation (Zeithamova et al., 2012; Preston & Eichenbaum, 2013). Several lines of evidence suggest that this area supports the neural implementation of a ramping-to-threshold decision process, in which accumulated evidence toward a choice is built over time until a response is provided (Hanks et al., 2015).

Characterizing such processes can benefit from the use of computational models that estimate latent variables - unobservable parameters that capture underlying cognitive mechanisms guiding behaviour. A typical framework is the drift diffusion model (DDM), which has been extensively studied in perceptual decision-making tasks. The DDM simulates decision processes as an overtime accumulation of evidence, which rate of accumulation may depend on the difficulty of the decision. In perceptual decision-making tasks, the drift rate is modulated by the degree of similarity of the perceptual stimuli to be evaluated (Gold & Shadlen, 2007; Ratcliff & Mckoon, 2008). A similar process likely underlies decision-making in TI, where the drift rate could vary as a function of the SDE - a similarity measure based on non-perceptual properties of the stimuli, with larger values yielding faster evidence accumulation (Ratcliff, 2022).

Within this framework, one could ask whether individual differences in PFC function may affect inferential reasoning performance. The Val158Met single-nucleotide polymorphism in the *Catechol-O-Methyltransferase* (*COMT*) gene is one element that contributes to such variability among individuals. It leads to a valine (Val) to methionine (Met) amino acid substitution, which affects the activity of the COMT enzyme at body temperature (Lachman M. et al., 1996). The Met allele is associated with lower COMT activity and, as a consequence, higher extracellular dopamine (DA) availability in the PFC compared to the Val allele, caused by the slower DA degradation enzymatic activity (Chen et al., 2004; Bilder et al., 2004). DA availability plays a critical role in modulating executive functions and cognitive control processes mediated by the PFC (Ott & Nieder, 2019). Accordingly, the COMT Val158Met polymorphism has been linked to individual differences in cognitive functions, including cognitive stability and flexibility (Dickinson & Elvevåg, 2009).

In the present study, we examined the performance of individuals with different *COMT* genotypes (Met/Met, Val/Met, Val/Val) on the TI task. We observed that, while the performance in the test phase across genotypes was comparable, during the learning phase, clear differences emerged, with genotypes varying in the number of learning blocks required to reach the imposed threshold in performance. Applying the DDM, we also observed that performance in the TI task was accounted for by variations in the drift rate.

## 2. Methods

### 2.1 Participants

Eighty-three healthy participants previously genotyped for the COMT Val158Met polymorphism (Mione et al., 2015) were recruited to participate in the study. All participants had normal or corrected-to-normal vision and no history of significant drug or alcohol abuse, head trauma, or significant medical illness in the past 6 months. Participation in the study was voluntary, and informed consent was obtained in accordance with the protocol approved by the ethics committee of the Department of Psychology of Sapienza University. The *COMT* genotyping resulted in 20 Met/Met (Male=6; Female =14; mean age: 25 ± 2), 24 Val/Val (Male=10; Female =14; mean age: 25 ± 3) and 39 Val/Met (Male=20; Female =19; mean age: 25 ± 3). Genotype frequencies, within each group, were consistent with Hardy-Weinberg equilibrium (**χ² =** 0.278, df = 1, *p* = 0.59).

### 2.2 Genotyping

First, genomic DNA was obtained from buccal swabs and extracted using the Invitrogen Pure Link Genomic DNA Mini kit (Invitrogen Life Technologies, Carlsbad, CA, USA; Cat # K1820-01), following the manufacturer’s protocol. To maintain privacy, DNA samples were anonymized upon collection. The SNP Val158Met of the *COMT* gene was genotyped using a TaqMan SNP Genotyping Assay (Applied Biosystems, CA, USA; Assay IDC_25746809_50, Lot Number 1607104). PCR amplification was conducted with 5–10 ng of genomic DNA, 1X TaqMan Genotyping master mix (Applied Biosystems), and allele probes labeled with 5’-VIC or 5’-FAM fluorophores. The amplification reaction conditions on an EcoTM Illumina thermal cycle were as follows: 1 cycle of 95°C for 10 minutes, followed by 50 cycles of 92°C for 15 seconds and 60°C for 1 minute. To verify the *COMT* genotype, the Val158Met polymorphism was also assayed using the restriction fragment length polymorphism method. The polymorphic region was amplified by polymerase chain reaction (PCR) with the following primers: COMT_For 5’ GGGGCCTACTGTGGCTACTC 3’ and COMT_Rev 5’ TTTTTCCAGGTCTGACAACG 3’. The Val and Met alleles were distinguished by digesting the PCR product with NlaIII restriction enzyme, followed by 2% agarose gel electrophoresis and visualization using ethidium bromide staining. The undigested PCR product (109 bp) carried the G variant (Val), while the digested product, yielding two fragments of 96 and 13 bp, respectively, contained the A allele (Met). To ensure the accuracy of SNP genotypes, multiple replicates, reference DNA samples, and negative controls lacking DNA were included in the analysis.

### 2.3 Transitive Inference Task

Participants completed a computerized TI task, following the experimental protocol developed in previous studies (Brunamonti et al., 2011; 2017). The task consisted of learning and testing. During the learning phase, participants were presented with all pairs of adjacent items from a rank ordered series (Figure 1A; A>B>C>D>E>F) composed of six abstract images (black and white bitmaps - 30 x 30 mm - each with the same proportion of white area over a black background) and asked to identify the higher-ranked item in each pair by trial-and-error.

**Figure 1.**
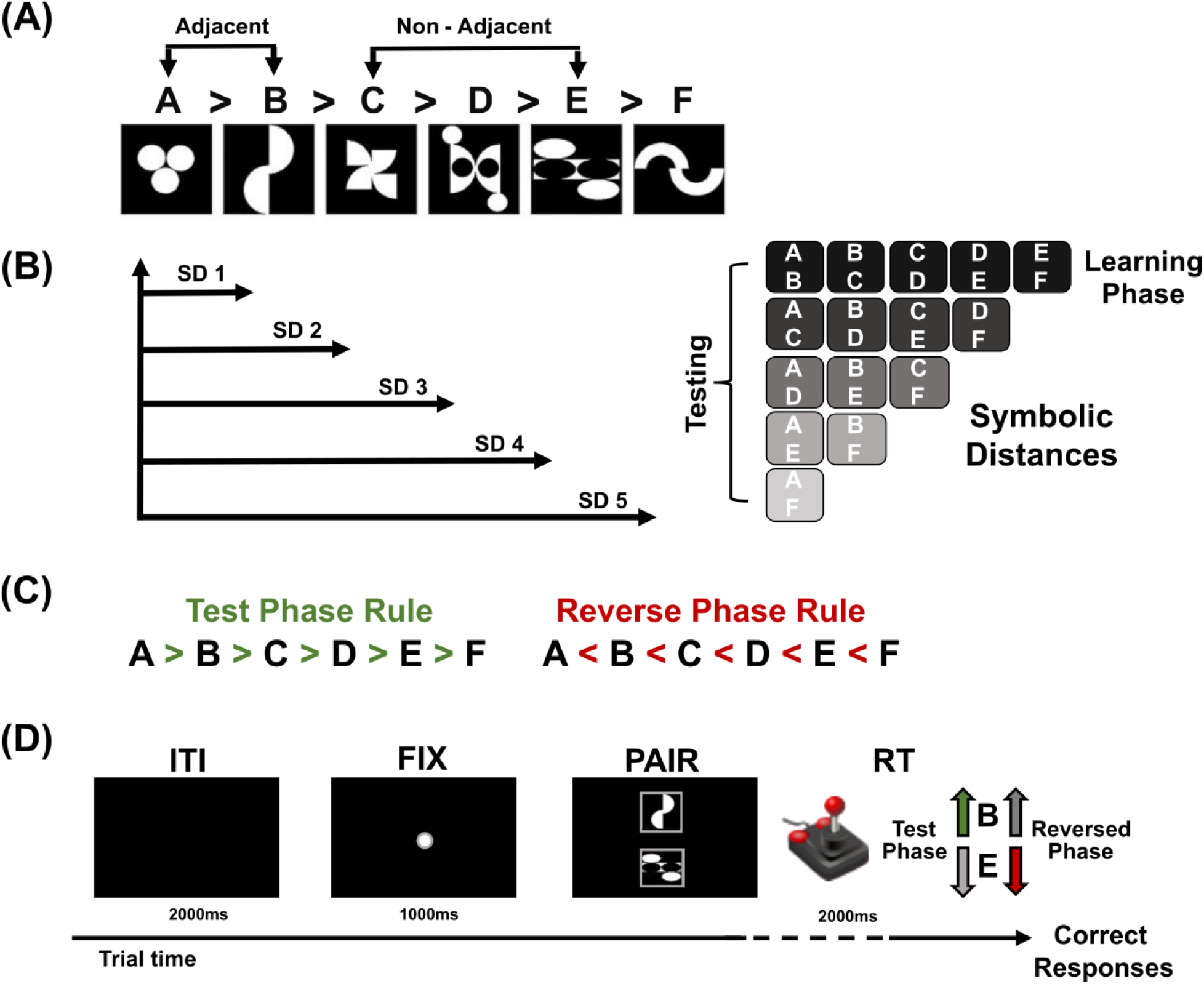
Transitive Inference (TI) task. **(A) Rank-ordered items**. Adjacent items for Learning and non-adjacent items for testing phase **(B) Symbolic Distances (SDs) and pair combinations**. During the testing, 15 pairs contributed to each SD, ranging from SD1 to SD5. Pairs were presented with the target item appearing randomly in either the upper or lower position. The panels on the left illustrate all the pair combinations from SD1 to SD5. (**C**) **Two testing Phases in this experimental design.** Test Phase rule: A>B to E>F and Reverse Phase rule: A<B to E<F. (**D**) **The sequence of events for each trial**.

All pairs of adjacent items were randomly presented in blocks of 50 trials (10 trials for each pair). Blocks of learning trials were presented until participants reached an average performance accuracy of 70%. At the end of each block, participants received feedback on the proportion of correct answers.

In the test phase, participants compared both learned and novel pairs that were never presented together during the learning phase (e.g., B>D or C>E; see Figure 1B for all pair combinations and SDs). During this phase, participants completed 150 trials, with each pair presented 10 times in random order.

To further investigate the relationship between the presumed difference on DA availability and the ability to adapt to new rules, we directed a subgroup of participants (48 out of 83) to engage in a second testing phase, a reverse phase, where the item’s rank order was reversed (see Figure 1C). Participants were informed about this reversal before starting the second testing phase. Each genotype group was divided into subgroups for the reverse phase, with 12 participants for Met/Met, 15 for Val/Val, and 21 for Val/Met.

Before starting, each participant received written instructions and was made familiar with the apparatus and the task. Participants were seated at a comfortable distance of 50 cm from a PC monitor in a dimly lit room. In both the learning and test phases, each trial began with a fixation point, a central target, displayed in the middle of the screen (see Figure 1D). After a delay of 1 second, a pair of items appeared on the screen, positioned one above the other. Participants used a joystick that was fixed to the table, aligned with their body midlines, and connected to a computer via USB, to indicate their choice by moving the joystick either forward or backward, as quickly as possible, to respond. If no response was made within 2 seconds of pair presentation (upper time limit), the trial was aborted and a new one began. Participants received two different auditory feedback indicating whether their response was correct or incorrect. Visual stimuli presentation and data acquisition were controlled by the freely available Psychtoolbox software (MATLAB-based routines; www.psychtoolbox.org).

### 2.4 Behavioural Data Analysis

To evaluate task performance related to decision difficulty, across COMT Genotypes (Met/Met, Val/Val, Val/Met) in both phases of the task (learning and test phases), we conducted two separate two-way mixed model ANOVAs on participants’ accuracy and reaction time (RT). In the test phase, we used SD (from 1 to 5) as a within-subjects factor, while in the learning phase, we used adjacent pairs (A>B, B>C, C>D, E>F, E>F) as a within-subject factor. Additionally, we tracked the number of learning blocks for each participant required to complete the learning phase by reaching the accuracy criterion (70% correct on all adjacent pairs). This measure was used to assess potential differences in learning across genotype groups. Chi-Square and Kruskal-Wallis tests were used to verify if differences in learning processes were statistically significant.

In the subgroup of 48 participants performing the reverse phase, to evaluate the participants’ ability to follow the new rule, we used a three-way mixed model ANOVA adding Task phases (Test/Reverse) as a within-subject factor to compare the performance between the two phases across the Genotype groups. Significant effects were followed up by post-hoc LSD tests. Reference values for effect size measurements, yet ***ηp^2^***, are as follows: 0.01 is the small effect size; 0.06 is the medium effect size; and 0.14 or higher is the large effect size. Analysis was performed by custom functions developed in MATLAB® R2022b (www.mathworks.com) and Statistica (StatSoft, Inc., 2004. STATISTICA (data analysis software system), version 7 (www.statsoft.com).

### 2.5 Drift Diffusion Model Analysis

To assess which latent variable could account for performance differences across SDs and between Genotype subject groups, we fitted the DDM to each participant’s distribution of RT data using the PyDDM package (Shinn et al., 2020), aiming to identify which parameters best explained decision dynamics in our task. The DDM conceptualizes decision-making as a ramping-to-threshold process where the time of deliberation depends on several parameters such as the starting point (***z***), the drift rate (***v***), the decision boundary (***a***), and the non-decision time (**ter**), as the time capturing sensory or motor processing unrelated to the decision (Ratcliff & Mckoon, 2008).

As in previous reports (Ratcliff & Mckoon, 2008; Gomez et al., 2007; Pirrone et al., 2017; Hadian Rasanan et al., 2021) in the present study, we assumed that item discriminability difficulty, indexed by SD, would influence the decision parameters without affecting the non-decision time. Here, we focused on the parameters ***v*** and ***a*** as variables influenced by decision difficulty. ***z*** was kept fixed since previous studies highlighted that this parameter is mainly influenced by motivation or reward expectation, which are variables not influencing the execution of the present task (Mulder et al., 2012); (Giuffrida et al., 2023). In addition, when fitting the test phase data, the model’s parameters were assumed to vary linearly with increasing symbolic distance, whereas during the learning phase, the parameters were assumed to follow a U-shaped pattern (Serial Position Effect), favouring the documented comparisons of extreme items over those of middle items (Jensen, 2017). The model analysis was performed by evaluating the goodness of fit of three different models: 1) a null-model, used as a control, in which no parameter varied across SDs; 2) a drift-only model, in which only the drift rate was allowed to change linearly with SD; and 3) a drift-boundary model, in which both drift rate and decision boundary were allowed to change linearly with SD.

The parameter ranges were defined a priori (drift rate ∈ [–5, 10]; boundary ∈ [0.2, 3]), and the best-fitting model for each participant was selected based on the lowest Bayesian Information Criterion (BIC; Raftery, 1995). A lower BIC value indicates a better fit, whereas higher BIC values suggest either a poorer fit or unnecessary model complexity. To statistically assess whether DDM parameters differed significantly, we first performed a two-way mixed-model ANOVA (Genotype x SD) and (Genotype x Adjacent Pairs), including all participants who completed the standard test phase and learning phase. Additionally, to examine whether parameters varied across task phases and genotypes, we conducted a three-way mixed-model ANOVA (Genotype x SD x Test Phase) on the subgroup of participants who completed both the test and reverse phases.

## 3. Results

### 3.1 *COMT* genotype impacts the progression of learning but does not impact performance in the test phase

First, we analysed whether, in the test phase, participants had properly acquired the reciprocal rank relationship between the six items by verifying if accuracy and RT were shaped by the SD and if this differed between subjects with different genotypes (Figure 2A). For this purpose, we performed a two-way mixed ANOVA with Genotype (Met/Met, Val/Met, Val/Val) and SD (from 1 to 5) as factors. The statistical test detected no significant main effect of Genotype (F_(2,80)_ = 2.63, p = .54, ηp² = .06), suggesting a comparable accuracy across groups, while there was a significant main effect of SD (F_(4,320)_ = 74,43, p < .001, ηp² = .48; post hoc p < .01), indicating that the rank relationship among items was not only acquired but this was significantly shaped by SD with increased accuracy at larger SDs. No significant interaction between the two factors was detected (F_(8,320)_ =.87, p = .54, ηp² =.021). Similarly, RTs were significantly modulated by the SDs (F_(4,320)_ = 233.32, p < .001, ηp² = .745; post hoc p < .001), with faster RTs for larger SDs, but no significant differences were found across Genotype groups (F_(2,80)_ = 1.21, p = .30, ηp² = .029) nor significant interaction (F_(8,320)_ = p = .75, ηp² = .018).

**Figure 2.**
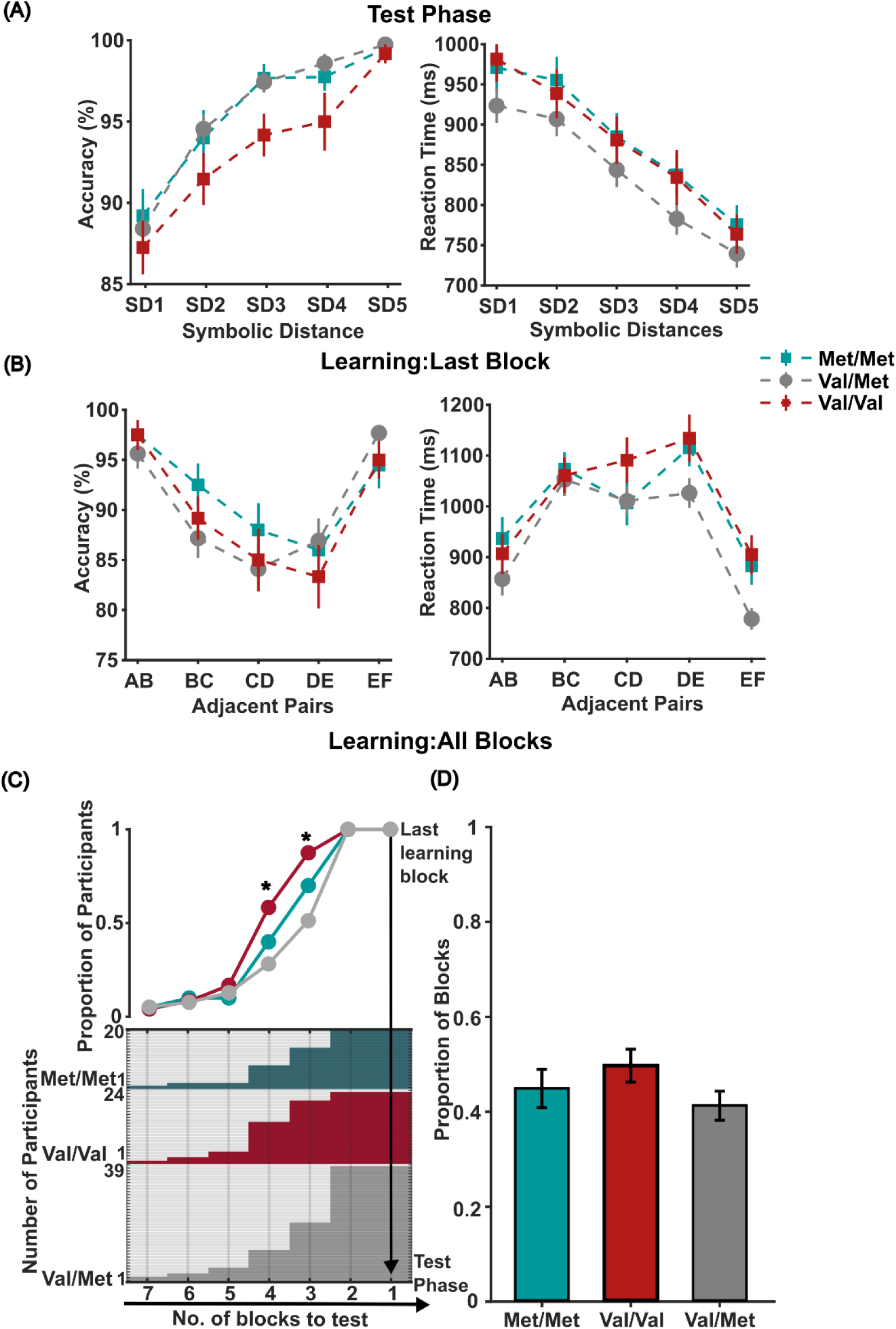
Genotypic COMT group performance in Test and Learning Phases. **(A)** Accuracy and Reaction Time as a function of SDs (from 1 to 5) for the entire Test Phase; **(B)** Accuracy and Reaction Time as a function of learned Adjacent Pairs for the last learning block; **(C)** Proportion of participants performing each learning block calculated from total learning blocks of a genotype group (top); Number of participants performing learning blocks across groups (bottom) **(D)** Proportion of learning blocks required by each genotype; Colored error bars in **(A)** are ±1 SEM among the pairs comprising each SD; in **(B)** are ±1 SEM among the pairs comprising each Adjacent pair. \****p*** < .05; respectively Met/Met (green squares), Val/Met (gray dots), and Val/Val (red squares).

Since the test phase revealed no significant genotype effects - meaning that, once the rank ordering was acquired, all groups performed alike - we redirected our analysis to the learning phase, especially focusing on how participants progressed across learning blocks. We first checked performance in the final learning block (Figure 2B) to assess whether any Genotype-related differences emerged. A two-way mixed ANOVA with Genotype (Met/Met, Val/Met, Val/Val) and Adjacent Pairs (A>B, B>C, C>D, E>F, E>F) was performed on accuracy and RT as factors. Results still showed no significant effect in accuracy for Genotype (F_(2,80)_ = 0.87, p = .42, ηp² = .021), nor a significant interaction (F_(8,320)_ = .77, p = .62, ηp² =.019). A significantly higher accuracy was found for Adjacent Pairs (F_(4,320)_ = 16.57, p < .001, ηp² = 0.171), with higher accuracy for the extreme pairs A>B and E>F compared to other pairs (post hoc p < .001). Similarly, RTs were significantly lower for the extreme item comparisons than middle items, Adjacent Pairs (F_(4,320)_ = 47.45, p < .001, ηp² = .372) and post-hoc p< .05. No significant differences between Genotype (F_(2,80)_ = 2,34, p = .10, ηp² = .055), or interaction was detected (F_(8,320)_ = 1.60, p =.123, ηp² =.038).

The former results highlight that all participants completed with comparable performance in the latest block of learning and the following test phase, with no difference between the different *COMT* genotypes. However, given the request to achieve a threshold level before engaging in the test phase, comparing the participants in the last learning block and the test phase might have plateaued any potential genotype-related variations occurring during the learning progression. To account for this possibility, we analysed the progression of each participant through the learning blocks. As a first step, we quantified the proportion of learning blocks needed by each group to achieve the accuracy threshold. Figure 2C (bottom) displays for each participant the number of learning blocks performed before the last block. Figure 2C (top plot) illustrates the proportion of participants of each group performing different blocks of learning. The rightmost proximal end of the line plot illustrates that all the participants needed at least two blocks before approaching the test phase, with no difference between genotypes. Similarly, the leftmost distal end displays that less than 1% of the participants required more than three blocks of learning, with no difference between the genotypes. The great majority of participants needed 3 to 4 blocks of learning. When comparing these proportions between the three groups, we observed a greater proportion of Val/Val individuals requiring 3 or more learning blocks and a lower proportion of Val/Met individuals. The Met/Met participants performed a proportion of learning blocks between the other groups. These differences were statistically significant for the last 3rd (χ²(2) = 10.85, p = .004) and last 4th (χ²(2) = 9.83, p = .007) blocks.

As a further control, we quantified for each participant the proportion of blocks required for achieving the set threshold, with respect to the maximum number of detected blocks across all the participants. Figure 2D displays the median of the distributions of these proportions, which was significantly higher in the Val/Val genotype, Kruskal-Wallis (χ²(2) = 6.57, p = .037). These findings indicate that while final block learning performance did not differ between genotypes, the learning process reflected potential genotype-related differences, with Val/Val participants requiring more trials/blocks to reach the same level of performance.

### 3.2 The difficulty level of the test phase modulates the drift rate underlying decision making

In searching to uncover variables underlying the decision process in the TI task, we modelled each participant’s performance by a DDM to evaluate which model’s parameters better accounted for observed behaviour.

Figure 3A displays the distribution of the reaction time of a representative participant for the five different levels of task difficulty, represented here by the SDE, and the simulated data by using the drift-only model, the drift-boundary model, and the null model. The evaluation of the goodness of fit of the model provided for 81 participants over a population of 83 (98%), the lowest BIC value for the drift-only model. In 1 participant, the lower BIC was associated with the null model, and in another one, the lower BIC was associated with the drift-boundary model (see Supplementary Figure 1 for details).

**Figure 3.**
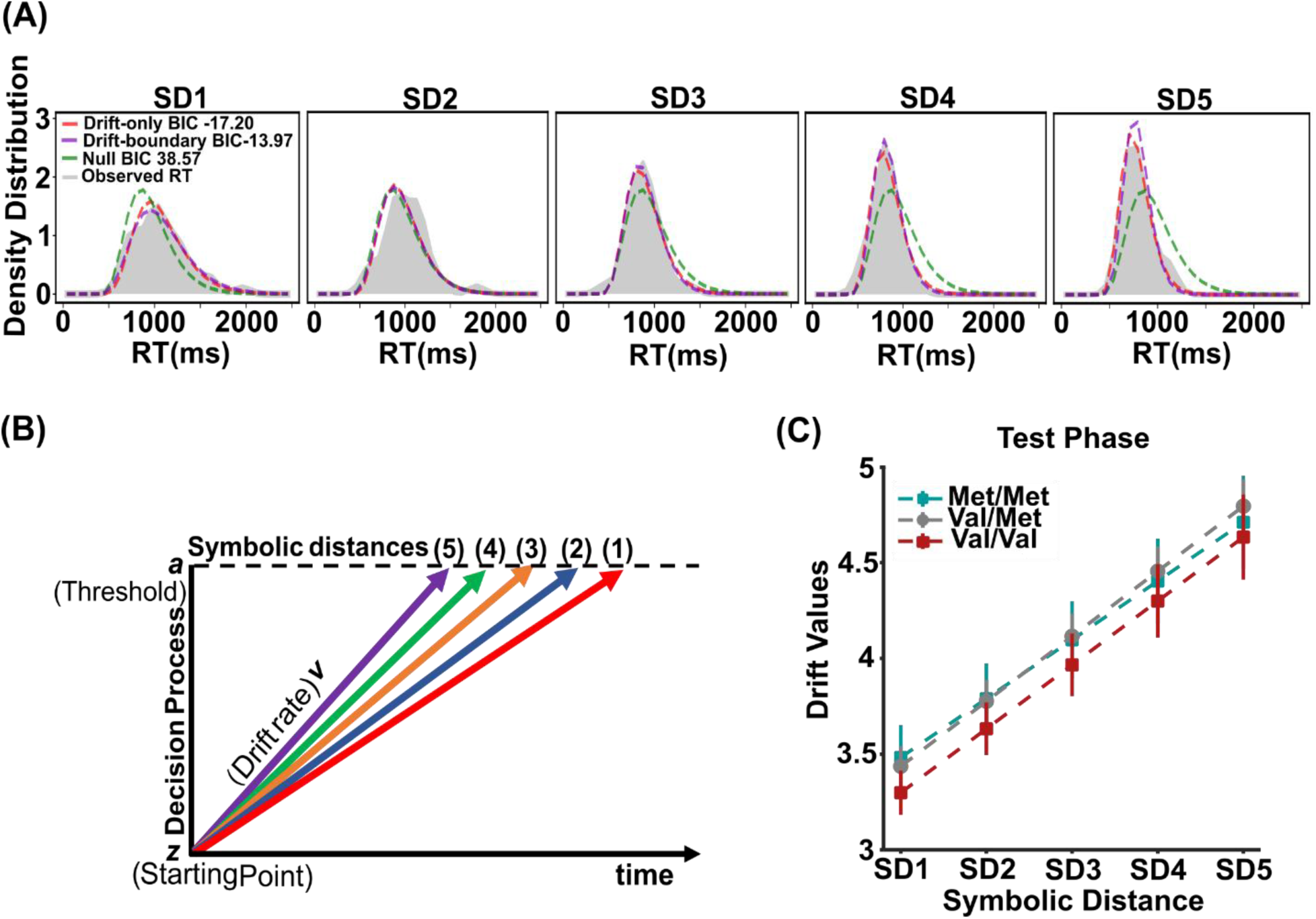
DDM simulation of RT distributions in the Test Phase. **(A)** Density distributions of reaction times for a representative participant across the five SDs, along with fits from the null model, drift-boundary model, and drift-only model. **(B)** Schematic representation of the DDM, where drift rate reflects the speed of evidence accumulation toward the threshold, across SDs. **(C)** Drift rates as a function of SDs (from 1 to 5) for the Test Phase. Coloured error bars are ±1 SEM among the pairs comprising each SD; respectively, Met/Met (green squares), Val/Met (gray dots), and Val/Val (red squares).

These results highlight the drift rate as the main variable varying with the task difficulty during the test phase. Consistent with the behavioural data, the drift rate differed significantly between the SDs, but it was comparable across the genotype groups (Figure 3C).

A two-way mixed ANOVA was conducted, with Genotype (Met/Met, Val/Met, Val/Val) and SDs as factors, we detected a significant main effect of SDs (F_(4,312)_ = .39, p < .001, ηp² = .848; post hoc p < .001) and a lack of statistical significance for the main factor Genotype F_(2,78)_ = .28, p = .75, ηp² = .007). No significant interaction was detected F_(8,312)_ = .38, p = .92, ηp² = .009.

### 3.3 Progressive learning of item relationships influences the decision drift rate

The previous results highlighted the drift rate as the main variable accounting for the difficulty-related performance in the test phase. With a similar approach to the test phase, we applied the DDM to assess if the drift rate accounts for performance from the start to the end of the learning phase. As in the test phase, in the last learning block, the drift rate was the variable that best accounted for task performance (Supplementary Figure 2). Figure 4A displays the different drift rates estimated by the model for the different pairs comparisons in the genotype groups in the last block of learning. The distribution of these values reflects the U-shape that shaped the reaction times (Figure 2B). A two-way mixed ANOVA was conducted, with Genotype (Met/Met, Val/Met, Val/Val) and Adjacent Pairs (A>B, B>C, C>D, D>E, E>F) as factors, detected a significant difference for Adjacent Pairs (F_(4,292)_ = 78.53, p < .001, ηp² = .518) while no significant differences in drift rates were observed across genotype groups (F_(2,73)_ = 2.46, p = .09, ηp² = .063). The analysis also revealed a significant interaction effect (F_(8,292)_ = 4.37, p < .001, ηp² = .107) (Figure 4A). Post-hoc LSD tests revealed highest drift values for extreme items comparisons (‘AB’ and ‘EF’) than middle items comparisons (‘BC’,‘CD’, ‘DE’), and a significantly higher drift for the ‘EF’ pair in the Val/Met genotype compared to other genotypes (post hoc p < .01). No other Adjacent Pair comparisons between genotypes reached significance.

**Figure 4.**
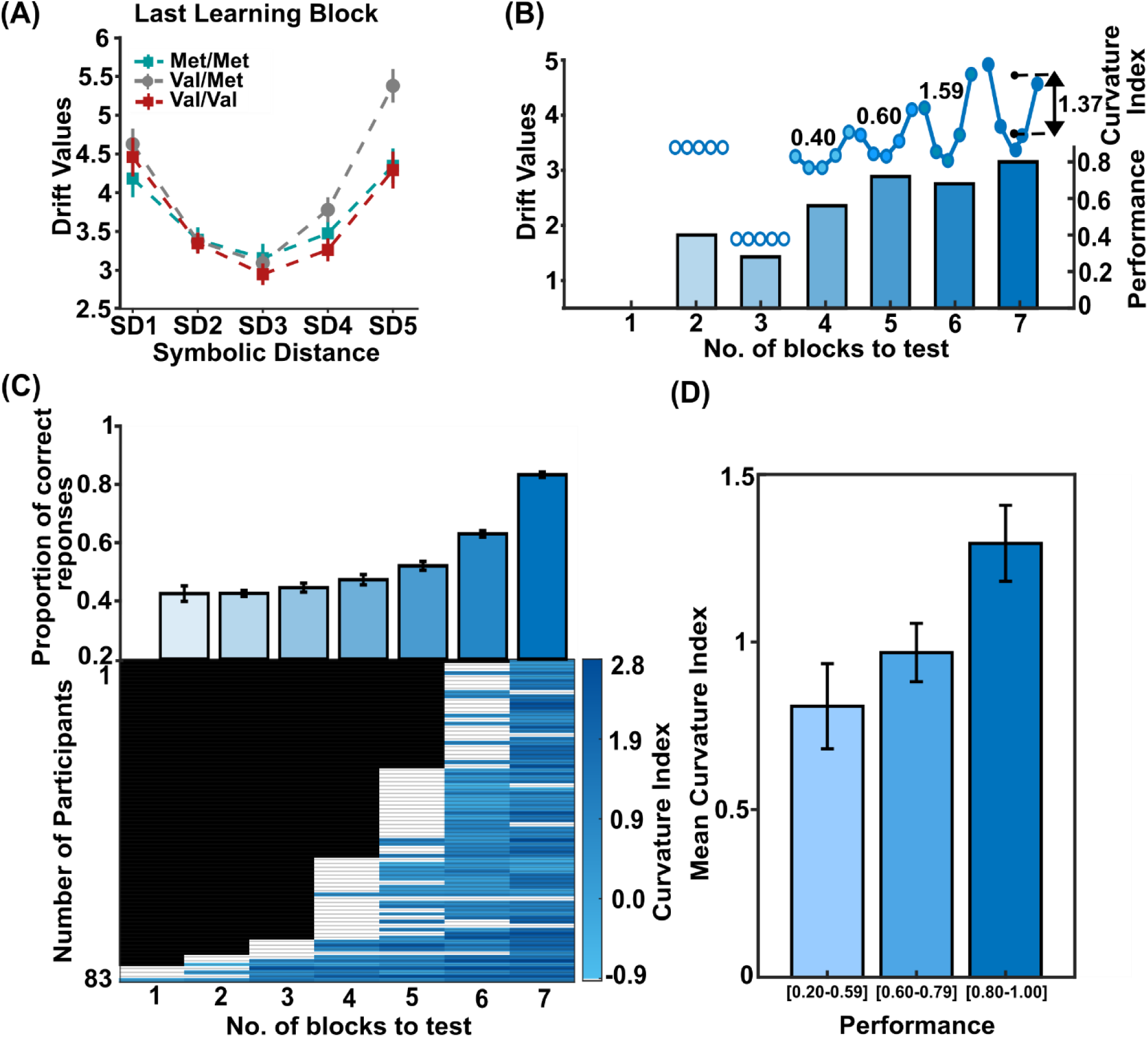
Performance and Drift rates in Learning Progression. **(A)** Drift rates as a function of learned Adjacent Pairs for the last learning block; Coloured error bars are ±1 SEM among the pairs comprising each Adjacent Pairs; respectively Met/Met (green squares), Val/Met (gray dots), and Val/Val (red squares) **(B)** Accuracy and drift rate progression learning blocks of a representative participant; inset values indicate the Curvature index **(C)** Average Proportion of correct responses for all participants irrespective of genotype (top); Curvature index calculated from drift rates for all participants for all learning blocks (bottom) **(D)** Relationship between curvature index and task performance.

We then tested the progression of learning, first to assess overall performance improvement across learning blocks, and second, to identify when, during the progression of learning, the drift rate reliably started to account for the decision performance. For doing so, we calculated the average accuracy of all participants regardless of genotype or pair, for each block of learning. Next, for each participant and block, we fitted the different versions of the DDM and determined when the drift-only model achieved a lower BIC than the null model. Figure 4B illustrates this process in a representative participant, showing that after two learning blocks, the drift-only model fits better than the null model. As learning progressed and performance improved, the degree of curvature of the U shape increased with the improvement of the performance. This U shape, shown in Figure 2B, suggests that the comparisons between the extreme items (A>B and E>F) were easier to perform than those between central items. For each participant and learning block, we computed a curvature index as the mean drift rate of extreme pairs (AB, EF) minus the mean drift rate of middle pairs (BC, CD, DE).

Assuming that learning progression facilitates comparisons between extreme item pairs (A>B and E>F), we expected this curvature index to increase over blocks. Figure 4C (top panel) shows the increasing performance of all participants with progression of learning blocks, while the bottom panel shows when, for each participant, the curvature index began to reflect the serial position effect. The drift parameter emerged as significant for 12% of the participants in the 4th pre-test block, increasing to 33% in the 3rd block, while for the great majority of participants, 69%, the drift rate accounted for the task performance in the 2nd pre-test block, regardless of genotype.

Across all participants and irrespective of genotypic differences, the curvature index increased consistently when the drift-only model provided a better fit than the null model (white cells), following the trend of accuracy improving. Figure 4C shows the relationship between curvature index and task performance, measured as the proportion of correct trials. Given that the curvature index may reflect underlying drift values that influence performance, participants were grouped based on their accuracy levels (0.20-0.59, 0.60-0.79, 0.80-1.00) to assess whether differences in curvature were associated with variations in accuracy (Figure 4D). A one-way ANOVA revealed a significant increase in the curvature index for the three different performance levels among the groups (F_(2,167)_ = 4.71, p < .010, ηp² = .054). Post hoc comparisons indicated significant differences in the curvature index between the highest accuracy level and the other two levels, suggesting this parameter is meaningfully associated with performance levels, regardless of the genotype.

### 3.4 The *COMT* genotype groups perform differently when the comparisons depend on a novel rule

The analysis of the learning progression (Figure 2C) detected that the Val/Val group needed more learning blocks to achieve the same performance as Met/Met and Val/Met before the test phase. During that learning phase, the participants were required to identify which items were reciprocally linked by the relational rule “higher than”. Then we implemented a variation of the task where we required a subgroup of participants to perform the same pairs of items’ comparisons, but with the reversed rank order (Figure 1C). Therefore, if the previously acquired sequence was A>B>C>D>E>F, in the new phase of the task, they were informed that the pair comparisons needed to rely on the sequence F>E>D>C>B>A.

Figure 5A illustrates the accuracy of the three genotype groups, both in the subgroup of participants completing the version of the task with respect to the initial version. The three way mixed ANOVA revealed significant effects for SDs (F_(4,180)_ = 100.26, p < .001, ηp² = .69), while no significant differences in Accuracy neither between groups based on genotypes (F_(2,45)_ = 1.83, p = .17, ηp² = .075) nor significant differences among Phases (F_(1,45)_ = .00, p = .95, ηp² = .00), were detected. Significant interactions between factors task phases and SDs (F_(4,180)_ = 2.74, p = .03, ηp² = .057) and all the three main factors were detected (F_(4,180)_ = 2.04, p = .04, ηp² = .083). The analysis of post-hoc identified a significantly lower performance in smaller SDs in the reverse phase (p < .05), which was mainly due to the lower performance of Val/Val (p < .05). No significant interaction was detected among task Phases and Genotypes (F_(2,45)_ = .76, p = .06, ηp² = .032).

**Figure 5.**
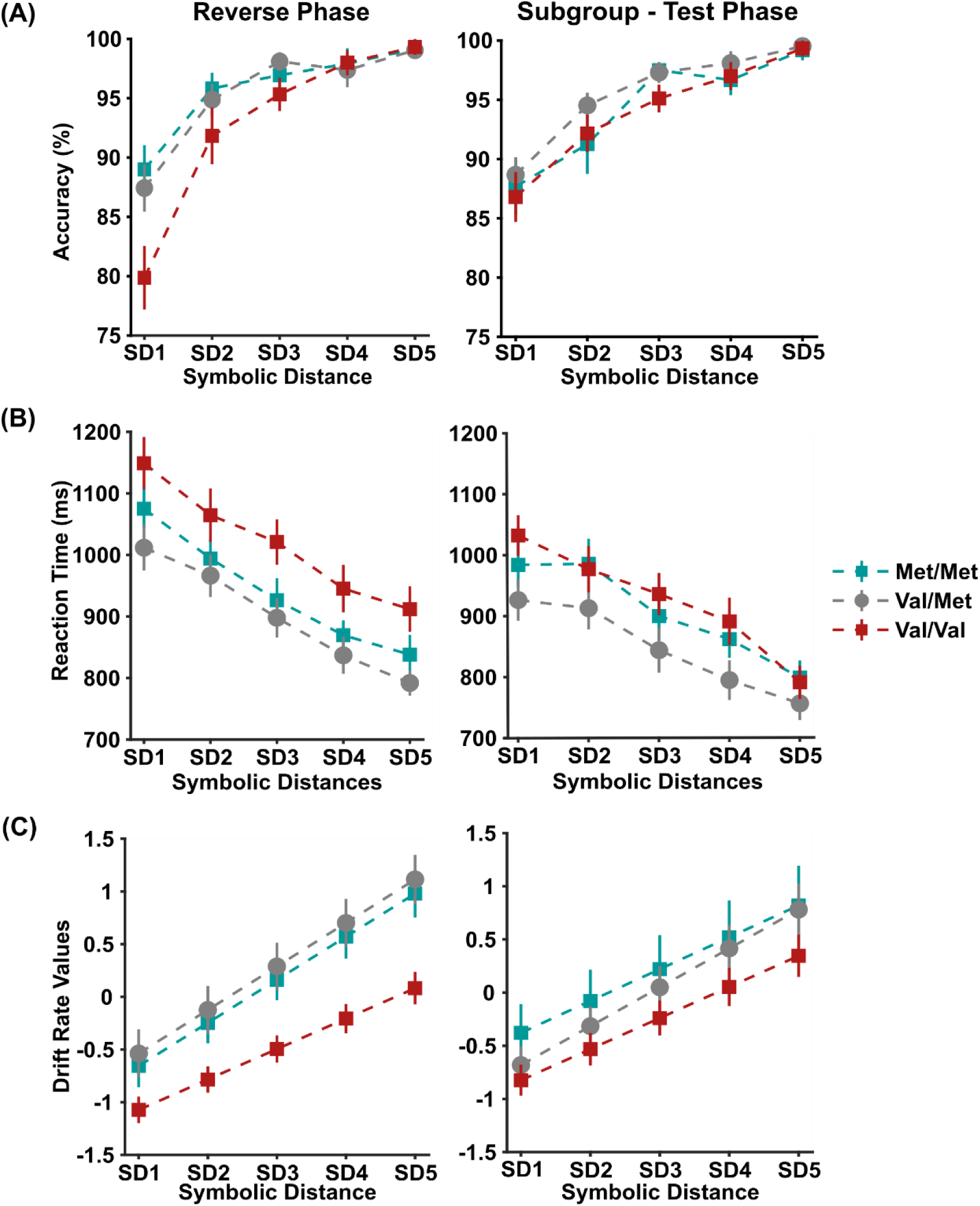
Genotypic COMT group performance and Drift rates in Reverse Phase. **(A)** and **(B)** Accuracy and Reaction Time as a function of SDs (from 1 to 5) for the Reverse Phase and Sub-group of Test Phase; **(C)** Normalized Drift rates as a function of SDs (from 1 to 5) for the Reverse Phase and Sub-group of Test Phase. Coloured error bars in **(A)**, **(B)**, and **(C)** are ±1 SEM among the pairs comprising each SD; respectively, Met/Met (green squares), Val/Met (gray dots), and Val/Val (red squares).

Different RT between genotype groups in the two task phases was also statistically significant. Figure 5B illustrates longer reaction times in the reverse phase than in the test phase. The three-way ANOVA revealed significant differences among the Phases (F_(1,45)_ = 12.65, p < .001, ηp² = 11.24). A significant effect of the SDs as a main factor (F_(4,180)_ = 219.36, p < .001, ηp² = .83) and an interaction of SDs with the task Phases was detected (F_(4,180)_ = 6.31, p < .001, ηp² = .051). Significantly different RT for SD1 and SD5 in the reverse phase were observed (post-hoc p<0.05). No significant effect of the main factor Genotype was observed (F_(2,45)_ = 2.97, p = .06, ηp² = .116), neither the interaction of genotypes with task phases (F_(2,45)_ = .94, p = .39, ηp² = .04) nor the interaction with SDs (F_(8,180)_ = 1.12, p = .35, ηp² = .047) was significant. No significant interaction among the three main factors was observed (F_(8,180)_ = 1.21, p = .29, ηp² = .051).

Even in the reverse phase, we used the DDM and detected the drift rate as the main variable accounting for the behavioural performance of each participant (Supplementary Figure 3). Prior to conducting the ANOVA, drift rates were normalized across participants to address substantial inter-individual variability and differences in the absolute drift rates between the Phases. Figure 5C illustrates the normalized drift rate obtained for the three genotype groups in the two task phases. The three-way mixed model ANOVA revealed significant differences among the genotypes (F_(2,43)_ = 3.44, p = .04, ηp² = .137). The drift rates of Val/Val were significantly lower than the other two groups post-hoc p < .05. Significant effects for SDs (F_(4,172)_ = 439.54, p< .001, ηp² = .001) and an interaction effect of SDs with Genotypes was detected (F_(8,172)_ = 3.42, p = .001, ηp² = .097), where Val/Val drift rate was smaller for SD4 and SD5 (post-hoc p < .05). The main factor Phases also had a significant interaction effect with SDs (F_(4,172)_ = 5.28, p <.001, ηp² = .00), with smaller drift rates for lower SDs in the reverse phase (post-hoc p < .05). There was no significant main effects for Phases (F_(1,43)_ = .028, p = .86, ηp² = .001) and neither its interaction with genotypes was detected (F_(2,43)_ = 1.06, p = .35, ηp² = .050). No significant interaction effects of the three main factors were observed (F_(8,172)_ =1.95, p = .055, ηp² = .098).

## 4. Discussion

### 4.1 Slower learning and rule-adaptation in the Val/Val genotype

In the present study, we investigated whether individual differences associated with variations in the *COMT* genotype affect the performance on a TI task.

The analysis detected a comparable ability among the three analysed genotypes in completing the test phase of the task, once a criterion threshold was achieved during the learning phase. This suggests that the genotype did not affect the inferential decision once the item’s relations were acquired.

The experimental approach was designed to detect differences in inferential decision making that reflect variations in PFC function linked to the *COMT* Val158Met polymorphism. This genetic variation modulates dopamine availability in the PFC, with Val/Val individuals showing the lowest levels due to higher COMT activity, Met/Met individuals the highest, and Val/Met intermediate (Lachman M. et al., 1996; Egan et al., 2001). Given the role of dopamine in supporting cognitive functions such as working memory, attention, and decision-making (Ott & Nieder, 2019; Friston et al., 2014), we hypothesized that the amount of time required to integrate the available information leading to a decision would reflect differences in dopaminergic tone across genotypes. Specifically, we expected Met carriers to exhibit higher accuracy and lower RT, while Val/Val individuals would show reduced performance due to lower dopamine availability. However, the lack of genotypic differences in completing the test phase contrasts with our initial hypothesis and with prior studies reporting lower cognitive performance in Val/Val individuals (Dickinson & Elvevåg, 2009).

Interestingly, we found that individuals with the Val/Val genotype required significantly more blocks of trials compared to Met/Met and Val/Met individuals to achieve the learning threshold, indicating greater difficulty in acquiring the items’ rank structure. A poorer performance of Val/Val also became evident in the reverse phase when participants were informed that the rank order of items was reversed.

Despite a comparable performance between *COMT* genotypes once they learned the items’ ranking, the performance diverged under conditions of reduced confidence in the relational hierarchy caused by the inversion of the original order of items. In this phase, Val/Val individuals exhibited reduced adaptability to the new rule. The lower performance of Val/Val in acquiring the rule of a new context seems consistent with the greater number of blocks needed by these individuals to complete the learning (see Figure 2C-D).

This might relate to the impaired reward-based associative learning of this genotype observed in other contexts (Lancaster et al., 2012; 2015). The Val/Val individual’s generally longer RT and lower accuracy for more challenging pair comparisons in the reverse phase are consistent with a diminished capacity of reviewing the learned hierarchical representation by associating positive feedback to the inverted pair association. This is probably due to the different amounts of dopamine available in the PFC. This interpretation aligns with previous research showing that Val/Val individuals performed poorly in a Stroop task with higher cognitive control demands, suggesting a common dopaminergic mechanism behind both rule updating and evidence evaluation (Rosa et al., 2010).

Together, these findings reveal a pattern in which individuals with the Val/Val genotype show reduced efficiency not only in the initial acquisition but also in the flexible updating of relational structures.

However, the present results contrast with previous studies reporting a cognitive flexibility advantage of Val/Val individuals, who exhibited lower switching costs in task-switching paradigms (Colzato et al., 2010; Moriguchi & Shinohara, 2018). This line of research emphasizes the Val allele’s role in rapid cognitive set reconfiguration (Markant et al., 2014), highlighting an aspect of flexibility that our data don’t seem to confirm. This reported discrepancy outlines the multifaceted nature of cognitive flexibility, which emerges in function of the task context. In our TI task, pair comparisons involve not only task switching but also the integration and restructuring of complex relational hierarchies, a task requiring deeper control processes. Our results suggest that Val/Val individuals would be at a disadvantage under these task demands since their lower dopamine availability hinders efficient and flexible adaptation, which is observed in demanding task-switching tasks (Rosa et al., 2010).

Future research should aim to disentangle the subprocesses that contribute to cognitive flexibility, such as set-shifting, hierarchical updating, and relational integration, across varying cognitive demands and in relation to the *COMT* genotype, to gain a clearer understanding of this complex neurocognitive construct.

### 4.2 DDM modelling

To gain a deeper understanding of the decision-making process going on in the present task, we investigated latent variables hypothesized to play a role in modulating decision processes using a DDM computational approach (Ratcliff & Mckoon, 2008). Here, we analysed two parameters of the model in relation to both the task difficulty and the group’s differences.

The drift rate represents the average speed of evidence accumulation towards a decision’s boundary (Ratcliff & Mckoon, 2008). This parameter has been used to characterize decision efficiency in perceptual decision tasks, where faster and more accurate decisions are associated with higher drift rates, driven in these tasks by more discriminable sensory inputs (Tavares et al., 2017; Beste et al., 2018). Like the drift rate, the other model’s parameters, as the decision boundary or the starting point of ramping, could influence the decision dynamics since they are sensitive to contextual factors. For instance, while perceptual difficulty mainly affects the drift rate (Gold & Shadlen, 2007; Bolam et al., 2024), motivational influences such as reward anticipation can shift the starting point of evidence accumulation (Giuffrida et al., 2023) or attention to accuracy modulate the decision boundary (Ratcliff & Rouder, 2000; Zhang & Rowe, 2014).

These parameters have also shown sensitivity to individual cognitive differences. For example, Ratcliff & McKoon, 2024 reported that age-related decline in drift rate mirrors reductions in the quality of mnemonic evidence. Similarly, Park & Starns, 2015 conducted a study using a numerosity comparison task, where participants had to judge which of two sets of dots contained more items. They found that the drift rate parameter was a reliable measure of numerical precision.

In clinical populations, DDM parameters can reveal compensatory mechanisms or processing differences. For instance, Pirrone et al., 2017 used a two-alternative forced-choice orientation discrimination test to examine perceptual decision-making in adults with autism spectrum disorder (ASD). In contrast to a reference stimulus, participants were asked to determine whether a visual stimulus was tilted clockwise or anticlockwise. They found that although ASD people responded more slowly, their accuracy was comparable to that of neurotypical participants. A further decision boundary, which indicates that participants with ASD were more careful, was the reason for the delayed replies. Drift rates did not differ between the groups, indicating that both people with and without ASD had comparable quality of perceptual information extraction. In contrast, children with ADHD have been found to exhibit slower drift rates, with no significant differences in decision boundary (Karalunas & Huang-Pollock, 2013). These studies demonstrated that the DDM is an effective tool for identifying hidden parameters controlling decision making across different tasks or group populations.

In the present work, we observed that the drift rate mainly accounts for the performance differences observed as a function of the acquisition of items’ rank relation, the comparison’s difficulty, and the group differences.

Firstly, here we provide evidence that the DDM framework could be extended to the TI task. As already observed in tasks based on perceptual discriminability, the same theoretical model could account for the accumulation of evidence by internal manipulation of learned relational information. We show that the drift rate may capture an underlying computational mechanism that seems to link both perceptual and non-perceptual decision-making. Here, the SD effect, where item pairs that are further apart in a hierarchy elicit faster and more accurate responses, was reflected in higher drift rates. This is likely due to easier access to more distinct relational representations (Lippl et al., 2024). In the present configuration of the task, we found that the drift rate was the main parameter influencing the performance of participants in the task.

Additionally, we observed that the drift rate value increases with increasing performance accuracy measured as both the general value across pair comparisons and between pair comparisons within learning blocks (Figure 4 C-D). The DDM thus revealed that the drift rate was the parameter most sensitive to task difficulty, also during the different blocks of the learning phase. These results align with previous observations that the same parameter is modulated by learning and practice, correlating with improved accuracy and decreased RT (Zhang & Rowe, 2014; Dutilh et al., 2009; Wagenmakers, 2009). This parameter also accounts for the lower behavioural performance of Val/Val in the reverse phase, which displays the lower drift rate value.

To conclude, the observed genotype performance relationship in the TI task has implications, helping in understanding the cognitive deficits observed in various neuropsychiatric and neurodegenerative disorders (Perkovic et al., 2017; Taylor, 2018; Fang et al., 2019). Conditions characterized by deregulated PFC dopamine, such as in schizophrenia, often present with impairments in relational reasoning and executive functions, including deficits in tasks akin to TI (Egan et al., 2001; Titone et al., 2004; Lopez-Garcia et al., 2013). Molecular genetic studies have consistently identified *COMT* polymorphism as a key candidate gene associated with ADHD symptoms (Sun et al., 2014; (Mizuno et al., 2017; Jung et al., 2019; Taylor, 2018). Recent neuroimaging studies further highlight this genetic link, showing *COMT* gene associations with decreased grey matter volume (Shimada et al., 2017) and abnormalities in cortical thickness and surface area in children with ADHD (Jung et al., 2019). Our findings suggest that the *COMT* genotype contributes to the variability and severity of these impairments, explaining the executive function deficits and difficulties in abstract thinking and hierarchical organization observed in individuals with conditions such as ADHD and schizophrenia (Brunamonti et al., 2017; Titone et al., 2004).

## CRediT authorship contribution statements

**Ann Paul:** Data curation, Formal analysis, Methodology, Visualization, Writing – original draft, Writing – review and editing. **Mariella Segreti:** Formal analysis, Methodology, Visualization, Writing – original draft, Writing – review and editing. **Isabel Beatrice Marc:** Formal analysis, Methodology, Visualization, Writing – original draft, Writing – review and editing. **Maria Teresa Fiorenza:** Methodology, Resources, Writing – review and editing. **Sonia Canterini:** Methodology, Resources, Writing – review and editing. **Surabhi Ramawat:** Writing – review and editing. **Giampiero Bardella:** Writing – review and editing. **Pierpaolo Pani:** Writing – review and editing. **Stefano Ferraina:** Conceptualization, Writing – review and editing. **Emiliano Brunamonti:** Conceptualization, Investigation, Formal analysis, Methodology, Project administration, Supervision, Visualization, Writing – original draft, Writing – review and editing.

