## Supplementary Materials for "Individual differences in learning and decision-making: the role of COMT Val158Met polymorphism in transitive inference"

### Supplementary Figure 1

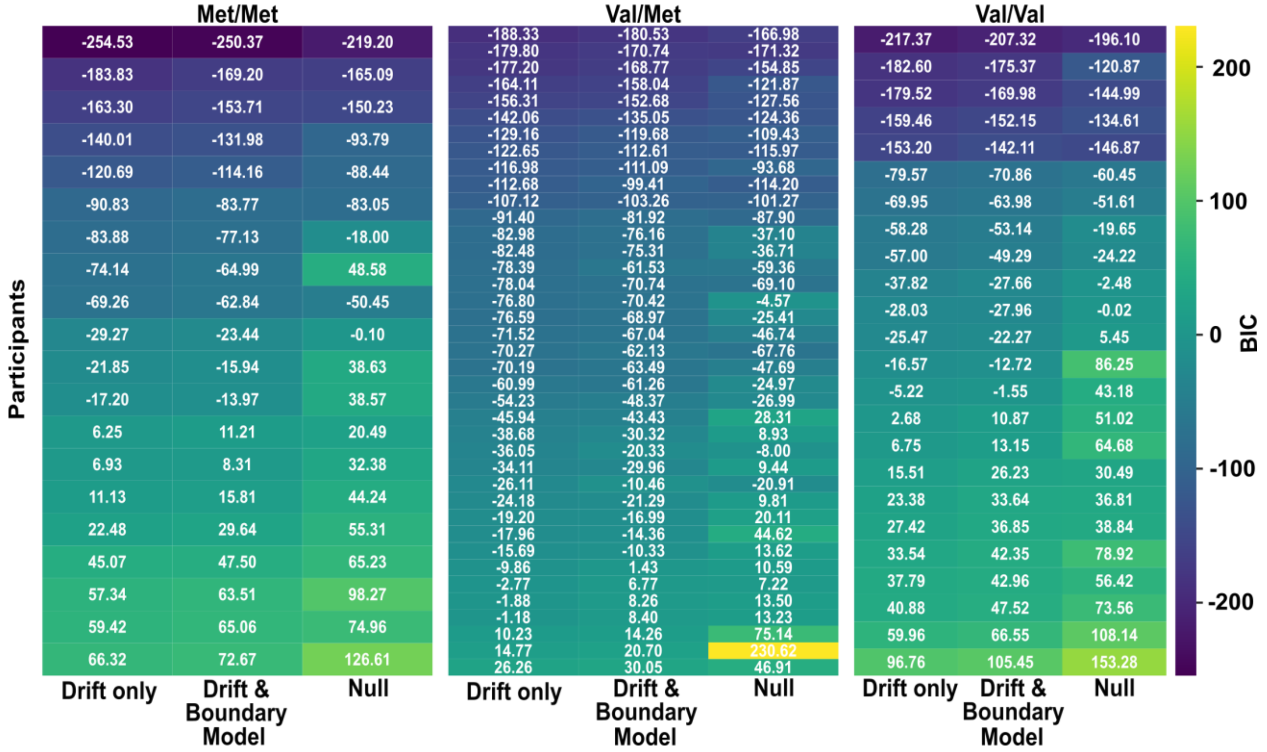


**Supplementary Figure 1**. **BIC Values of Test Phase for each participant** The Figure lists, for every genotype and participant, the BIC obtained when fitting three alternative DDMs to the RT distributions recorded in the test phase: (i) a **Null model** in which no decision parameter varies with SD, (ii) a **Drift-only model** allowing the drift-rate to scale linearly with SD, and (iii) a **Drift-Boundary model** in which both drift-rate and decision boundary vary with SD. Cells are colour-coded with a blue-yellow gradient in which the darkest color marks the lowest (best) BIC within each row.

### Supplementary Figure 2

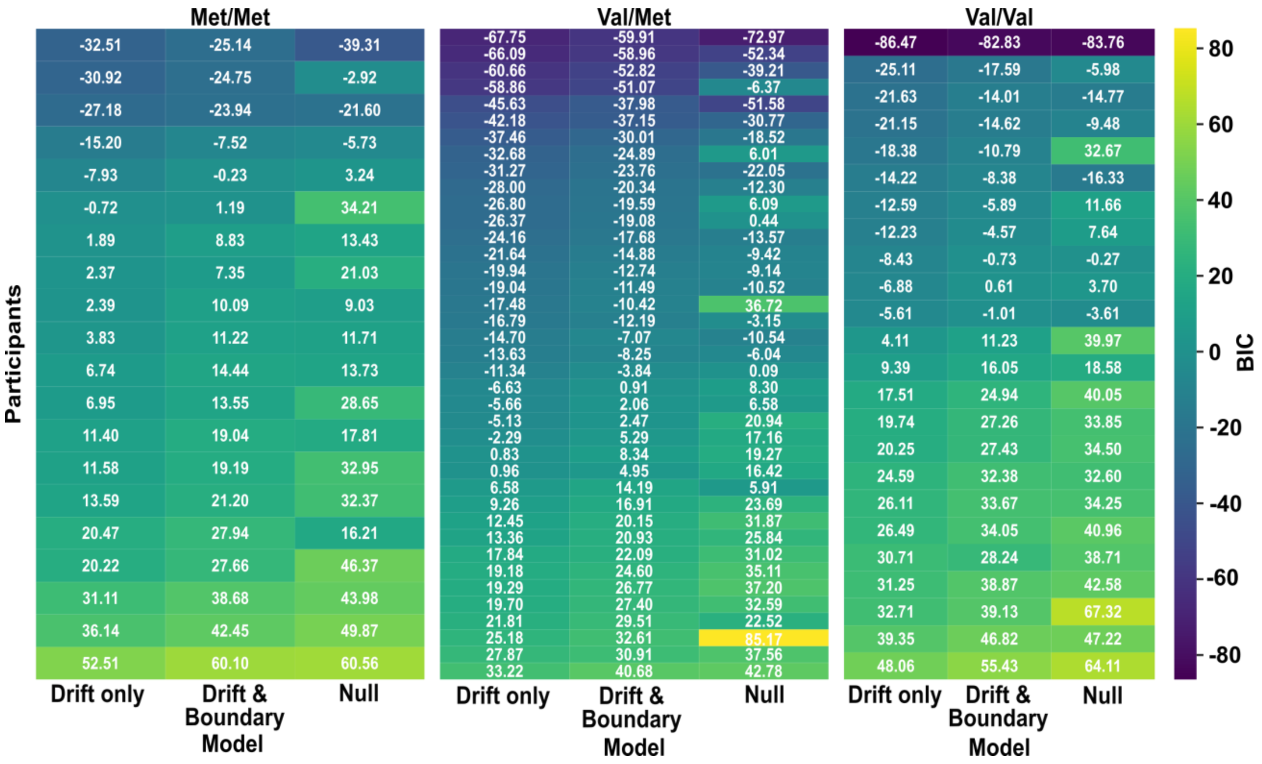


**Supplementary Figure 2**. **BIC Values of the Last Learning Block for each participant.** The Figure lists, for every genotype and participant, the BIC obtained when fitting three alternative DDMs to the RT distributions recorded in the final learning block: (i) a **Null model** in which no decision parameter varies with the pair, (ii) a **Drift-only model** allowing the drift to change with the pair, reflecting a U shape, and (iii) a **Drift-Boundary model** in which both drift-rate and decision boundary vary with the pair. Cells are colour-coded with a blue-yellow gradient in which the darkest color marks the lowest (best) BIC within each row.

### Supplementary Figure 3

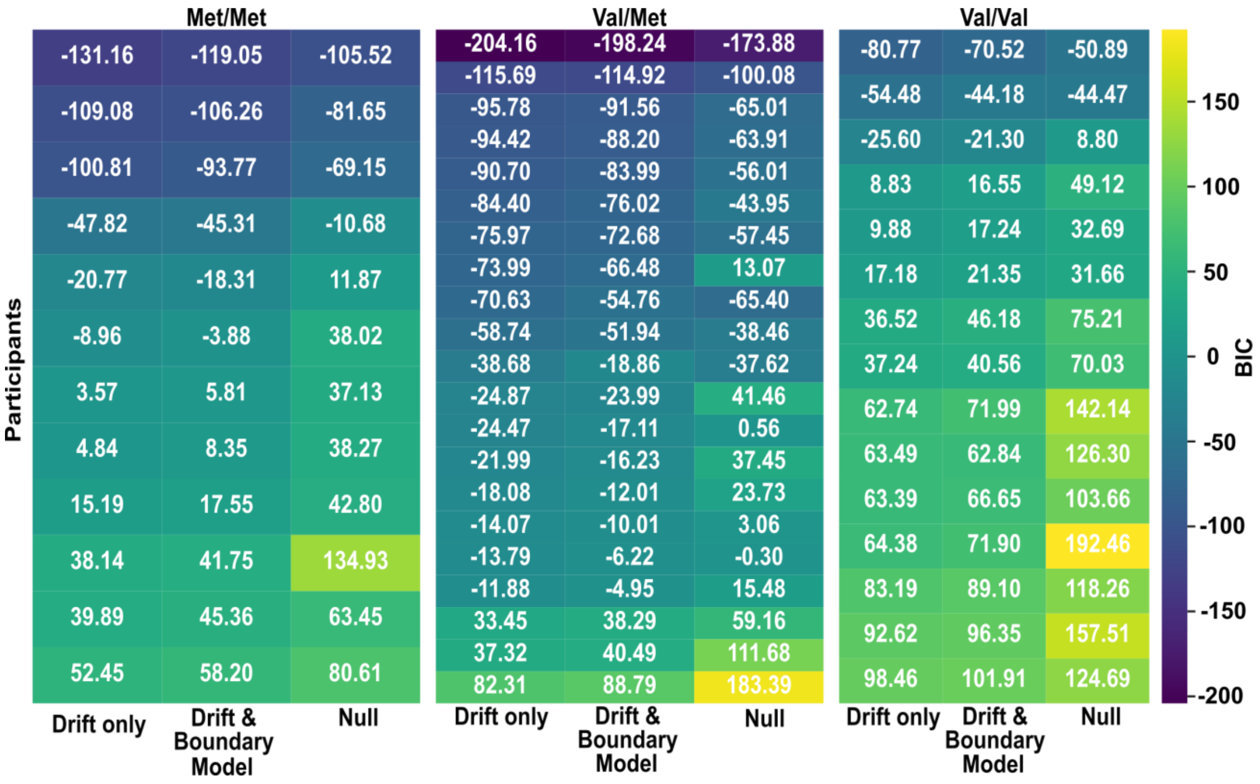


**Supplementary Figure 3**.**BIC Values of Reverse Phase for each participant.** The Figure lists, for every genotype and participant, the BIC obtained when fitting three alternative DDMs to the RT distributions recorded in the reversed phase: (i) a **Null model** in which no decision parameter varies with SD, (ii) a **Drift-only model** allowing the drift-rate to scale linearly with SD, and (iii) a **Drift-Boundary model** in which both drift-rate and decision boundary vary with SD. Cells are colour-coded with a blue-yellow gradient in which the darkest color marks the lowest (best) BIC within each row.
